# High-dimensional confounding in causal mediation: a comparison study of double machine learning and regularized partial correlation network

**DOI:** 10.1101/2024.10.12.617110

**Authors:** Ming Chen, Tanya T. Nguyen, Jinyuan Liu

## Abstract

In causal mediation analyses, of interest are the direct or indirect pathways from exposure to an outcome variable. For observation studies, massive baseline characteristics are collected as potential confounders to mitigate selection bias, possibly approaching or exceeding the sample size. Accordingly, flexible machine learning approaches are promising in filtering a subset of relevant confounders, along with estimation using the efficient influence function to avoid overfitting. Among various confounding selection strategies, two attract growing attention. One is the popular debiased, or double machine learning (DML), and another is the penalized partial correlation via fitting a Gaussian graphical network model between the confounders and the response variable. Nonetheless, for causal mediation analyses when encountering high-dimensional confounders, there is a gap in determining the best strategy for confounding selection. Therefore, we exemplify a motivating study on the human microbiome, where the dimensions of mediator and confounders approach or exceed the sample size to compare possible combinations of confounding selection methods. By deriving the multiply robust causal direct and indirect effects across various hypotheses, our comprehensive illustrations offer methodological implications on how the confounding selection impacts the final causal target parameter estimation while generating causality insights in demystifying the “gut-brain axis”. Our results highlighted the practicality and necessity of the discussed methods, which not only guide real-world applications for practitioners but also motivate future advancements for this crucial topic in the era of big data.

## 1 Introduction

Increasingly, causal mediation analyses are recognized by various domains. Upon evaluating the total causal effect of some treatment or exposure, investigators are further interested in its direct or indirect pathways through some mediator variable [15, 35]. For ubiquitous observational studies, the utmost challenge is to mitigate the selection bias by collecting as many baseline characteristics as possible. However, when the number of confounders approaches or exceeds the sample size, the model fitting becomes unstable without some filtering [4, 5]. In quantifying the causal mediation effect under high-dimensional confounders, selecting a subset of relevant confounders is essential to avoiding overfitting and drawing consistent inferential conclusions [41].

However, there is a gap in determining the best strategy for confounding selection in this growing problem of high-dimensional causal mediation analysis. In this paper, we focus on two promising approaches. One is the mediation extension of the double machine learning (DML) that incorporates penalization in modeling the nuisance functions that appeared in the efficient influence functions to handle the massive confounders [10]. Another is motivated by the emerging partial correlation network in deciphering the complex interplay among psychosocial variables [9]. Unlike the conventional Lasso [36, 42], which penalizes the coefficients in the regression model, the penalization is applied at the outset when fitting a Gaussian graphical model between the confounders and the response variable [17]. The deduced penalized partial correlation coefficients encourage selecting confounders with direct effects on the response while discouraging the selection of irrelevant ones by partial them out [41].

To address the gap and guide real-world applications, we illustrated different combinations of confounding selection methods to demystify the causality in the “gut-brain axis” in a motivating observational study [19, 30]. Our comprehensive comparison results offer methodological implications on how the confounding selection impacts the final causal target parameter estimation, above and beyond the valuable real-world scientific insights.

Essentially, our results are precious for the ubiquitous observational data, where the unknown confounding mechanisms are modeled as a nuisance using flexible machine learning approaches, especially when encountering massive confounders. The rest of the paper is organized as follows. In Section 2, we provide an overview of the motivating study on the gut-brain axis. In Section 3, we first detail the efficient influence functions (EIF) under the potential outcome framework in causal and mediation settings and then discuss the two confounding selection strategies. In Section 4, we offer comparisons among various combinations to evaluate their impacts on the final estimation of target parameters, which are applied to the motivating real-world data. We conclude the paper in Section 5.

## 2 Motivation

### 2.1 Background

The human microbiome consists of the microbes, their genetic elements, and their interactions with surrounding environments throughout the human body [7]. Numerous studies have suggested that the microbiome is the missing link between genetics, environment, and disease [21, 30, 38], incentivizing statistical advancements to decipher their inherent mechanisms, especially for the causal pathways, direct or indirect.

Fueled by the technological breakthrough of next-generation sequencing, the human microbiome composition can be interrogated using high-throughput sequencing. Marker genes can be amplified and sequenced and then clustered into Operational Taxonomic Units (OTUs) or amplicon sequence variants (ASVs). By comparing them with reference databases, one can establish taxonomic abundance profiles for each subject as the basis for statistical analyses [19]. Such abundance data are challenging to analyze. First, the number of taxa features exceeds the number of subjects in many studies. Second, those counts are quite sparse with a preponderance of zeros. Third, to overcome the heterogeneity or artifact in sampling depth, the absolute abundance is frequently normalized into relative abundance using centered log-ratio transformation, creating the compositional data that needs special statistical consideration. Lastly, the human microbiome is highly dynamic and varies on a day-to-day basis. Longitudinal studies with repeated measures will help depict more comprehensive insights but appropriate associational and causal inference tools are still lacking.

### 2.2 Gut-brain-axis

The gut-brain axis refers to the complex communications between the gut microbiota and the brain. For almost a century the human microbiota has been linked to neuropsychiatric disorders associated with neurodevelopment (e.g., autism spectrum disorder and schizophrenia), neurodegeneration (e.g., Parkinson’s disease, Alzheimer’s disease, and multiple sclerosis), and mood (e.g., depression and anxiety) [26]. However, we are still at the beginning of deciphering their innate mechanisms and causal pathways. A growing body of literature has suggested a strong connection between the gut microbiome and the central nervous system, evidencing the key role of the gut-brain axis. For example, in a cross-sectional study by **(author?)** [25], the beta-diversity, a measure of gut microbial community composition, was significantly associated with all measures of cognitive function. Also, major depressive disorder, a maladaptive response to chronic stress (or stress during early life), has been hypothesized to be mediated by the gut microbiome since s tress is a disruptor of gut microbiota composition in animals and humans [11]. Further, mechanisms underlying microbial-mediated changes in social behavior in mouse models of autism spectrum disorder have been confirmed [34]. Still, more causality evidence in human studies is needed to move beyond correlation to the validation of causal mechanisms.

### 2.3 Motivating study

Given most human gut-brain-axis research is limited by its observational and cross-sectional nature, showing association but not causation, a recent longitudinal study was conducted for a group of aging residents to overcome this challenge. In this study, demographics, physical, cognitive, and psychosocial instruments were assessed at baseline. Their fecal microbiome was sequenced at baseline and the follow-up visit after six months. Albeit observational, various potential confounders were collected (*p* = 81), which approaches the sample size (*n* = 92). Along with the time lag, it allows for repeated measures from the same individual to strengthen the cause-and-effect insights.

We hypothesize that exposure to loneliness and cognitively stimulating activities alters cognitive functioning over six months via the microbiome composition, which serves as the mediator (see Figure 1). The implication is pivotal in supporting the potential for gut microbiota targets in preventing or treating cognitive decline.

**Figure 1.**
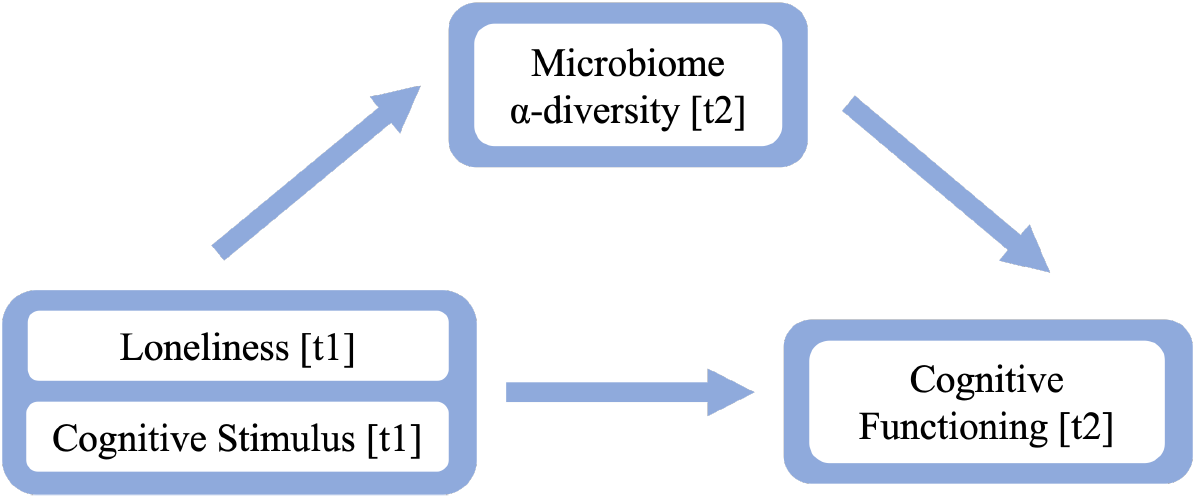
Hypothesized Causal Effect between Gut Microbiota and Cognitive Functioning

**Figure 2.**
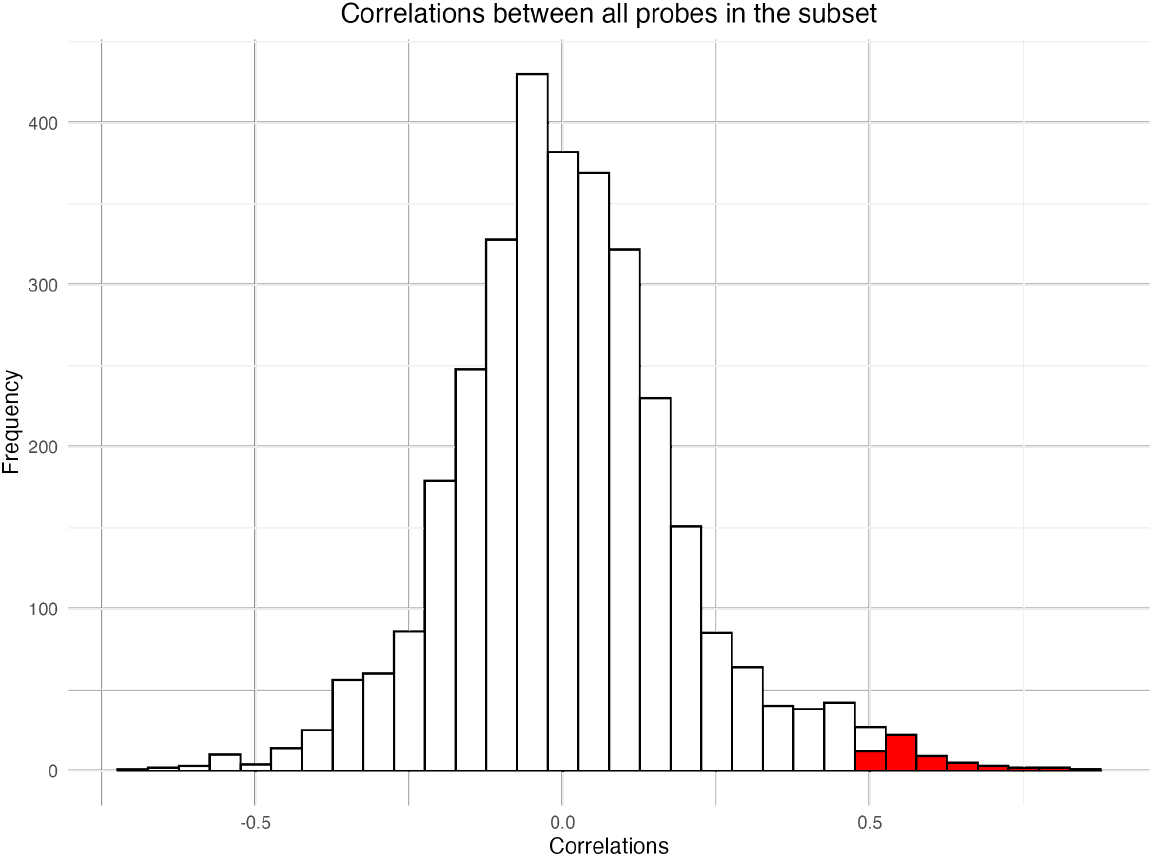
Pairwise Correlation Among Confounders

**Figure 3.**
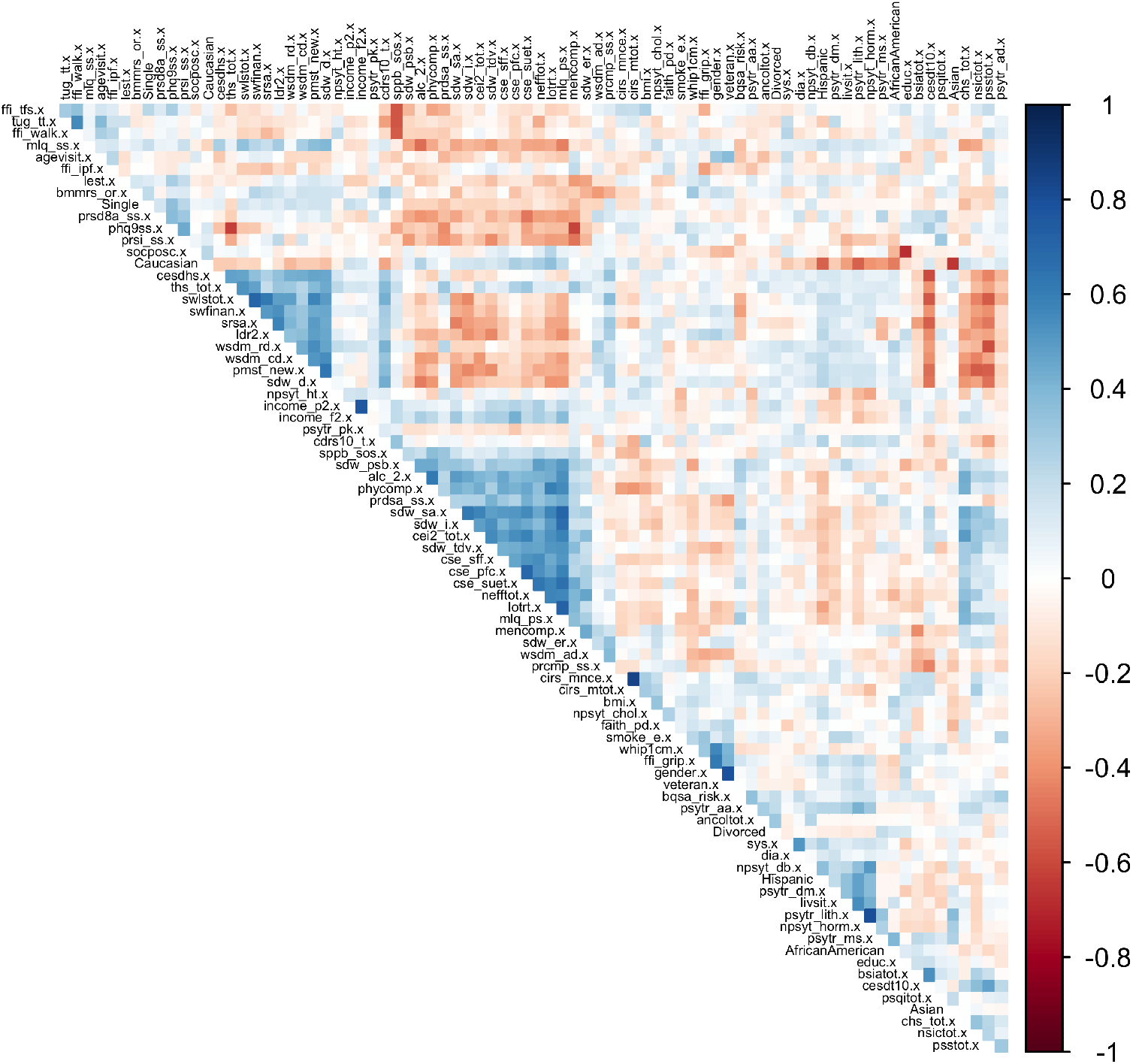
Correlation Heatmap Between Confounders

**Figure 4.**
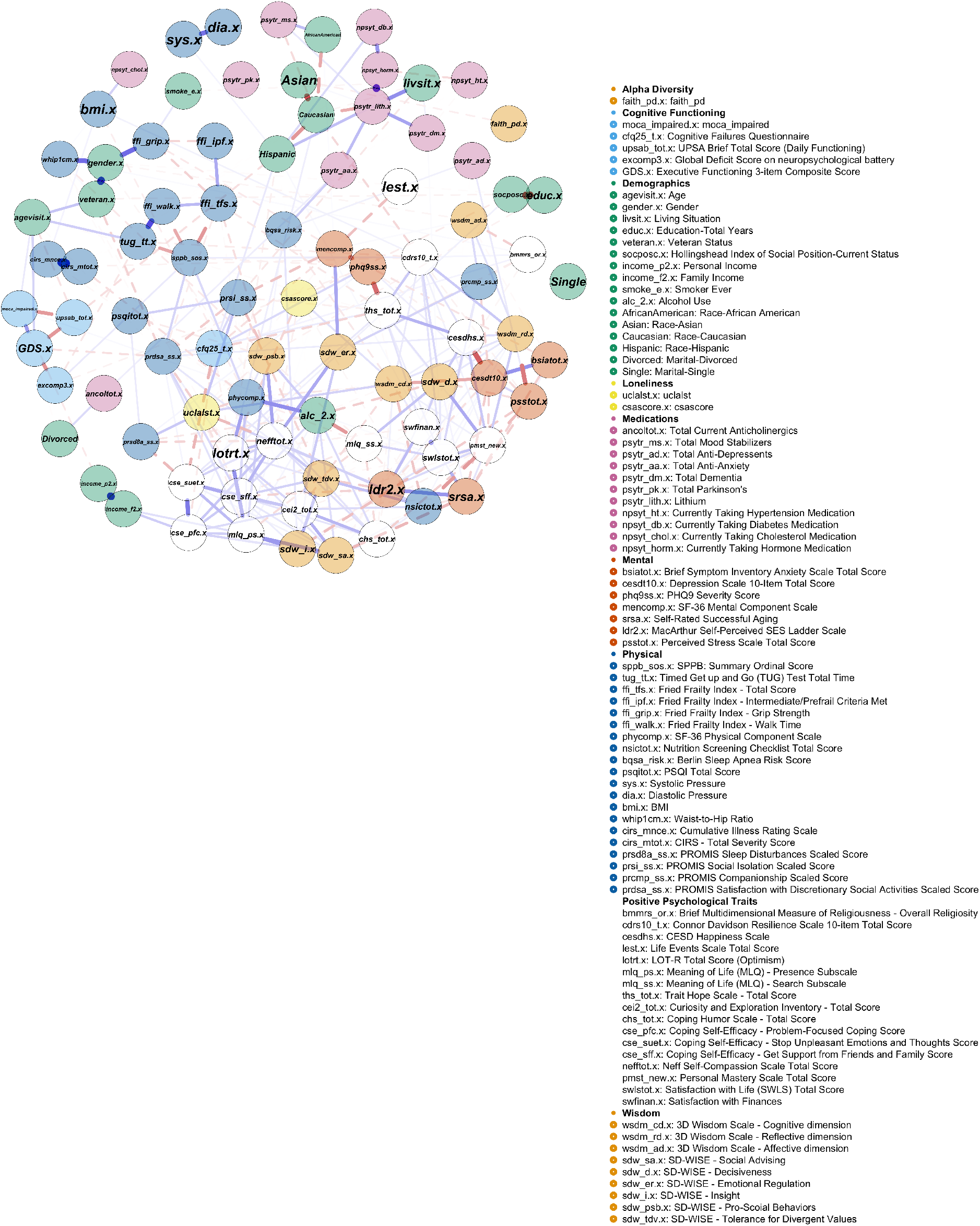
Network at *γ*=0

**Figure 5.**
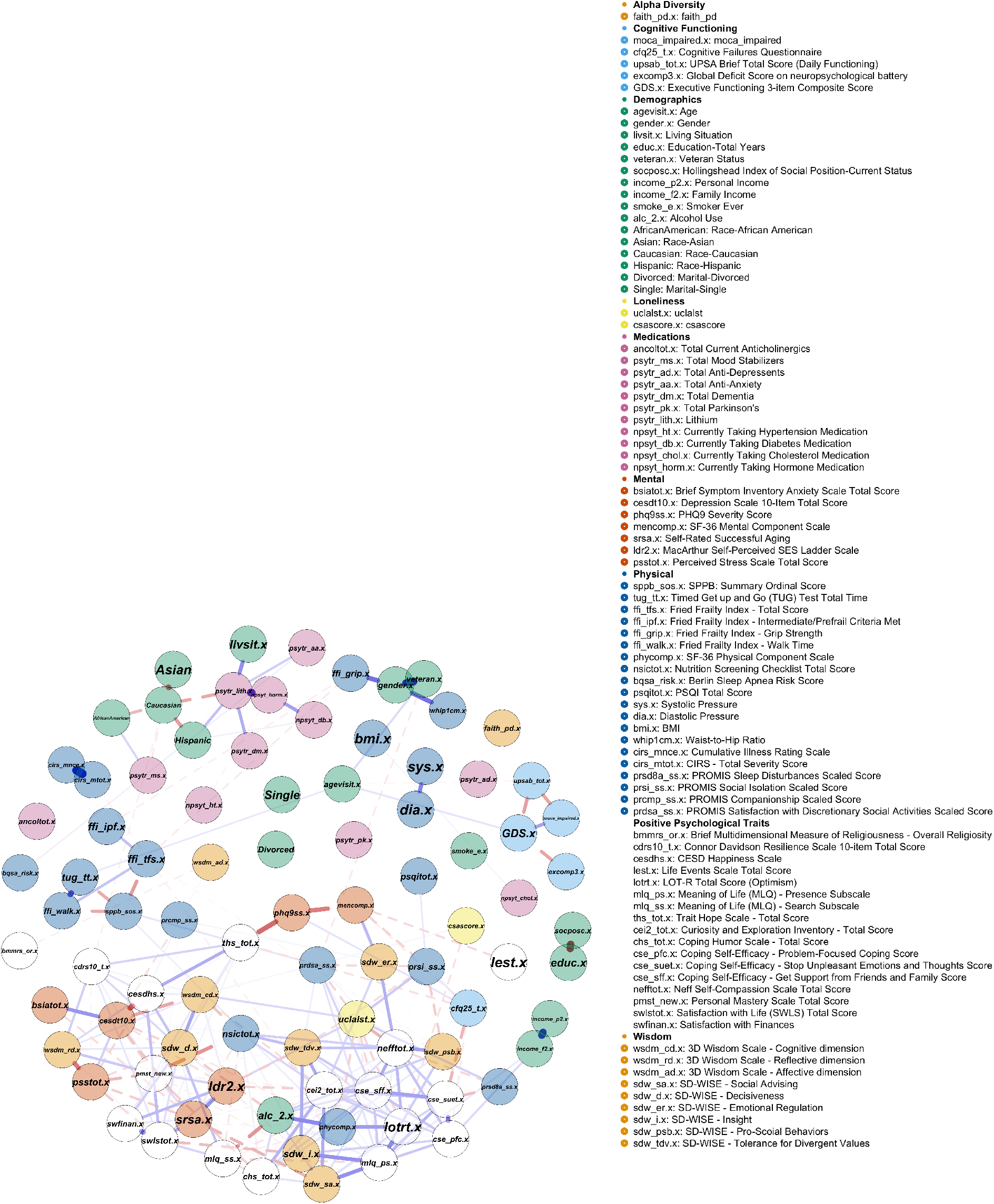
Network at *γ*=0.1

Denoted by **Y**_*i*_ = (*Y*_*i*1,_*Y*_*i*2,_…, *Y*_*ih*_) the microbial taxa counts at six months, compounding the issue of *n << h* = 363, **Y**_*i*_ are highly-skewed and zero-inflated as we mentioned earlier. An exploratory screening using the Lasso [42] indicated that 71 out of the 363 features had non-zero coefficients regarding cognitive decline. Investigating causal mediation impacts on the individual taxa based on this information will not only violate the rule of postselection inference [22] but is unlikely to survive the multiple comparisons, and hence, harm the study reproducibility. Whereas the feature aggregation provides a promising solution, specifically, we aggregated the taxonomic abundances at the within-subject level, yielding Faith’s phylogenetic alpha-diversity *f* (**Y**_*i*_), which also incorporates a biologically relevant tree structure across microbial taxa units [30]. Our goal is to investigate the modulation effect of the microbiome alpha-diversity on cognitive decline. Accordingly, the causal mediation pathway is examined by leveraging the exposure at baseline (loneliness, cognitive stimulating activities), the microbiome alpha-diversity as the mediator, and the cognitive functioning at six months as the outcome. We consider baseline measures of demographics, physical, and psychosocial instruments as potential confounders. To overcome the inferential challenge of massive confounders *n << p*, we focus on data-adaptive approaches such as double machine learning (DML) and regularized partial correlation networks to select a subset for further causality analyses.

## 3 Method

We review Neyman’s potential outcome framework in causal and mediation analysis.

### 3.1 Causal Effect

Consider a binary exposure *E*_*i*_ = *e* ∈ {0, 1}, where each subject is equipped with a pair of potential, or counterfactual, outcomes 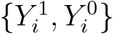 but only one of them is observed. The mean-difference type of *average causal effect* (ACE) aims to quantify the deviation of the outcome *Y*_*i*_ ∈ ℝ from the exposure arm to the control, averaging over the confounders **X**_*i*_ ∈ *χ* across the population [33]:

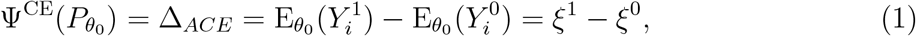

where the exposure-specific mean potential outcome 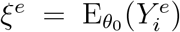 is termed the *causal functional*. Here 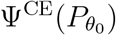 is identifiable with three assumptions:

C1. *SUTVA*: the observed outcome satisfies 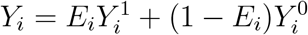.

C2. *Strong ignorability* :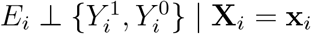.

C3. *Positivity*: Pr(*E*_*i*_ = *e* | **X**_*i*_) *>* 0, w.p.1. for each *e* ∈ {0, 1}.

To estimate and make inference about 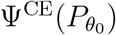, the *efficient influence function* (EIF) for *ξ*^*e*^ has been proposed [5, 37]:

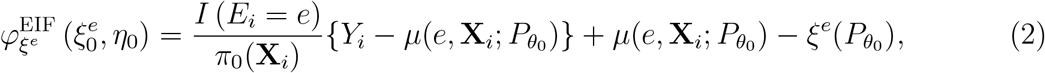

which is verified to be *Neyman orthogonal* [4, 29], and hence, the estimators of *ξ*^*e*^ solved from the estimating equations constructed from (2) are deemed doubly robust, namely, the final estimator is consistent, provide one of the two nuisance functions in (2) are specified correctly. The nuisance functions are the mean function 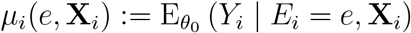 and propensity score (PS) 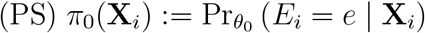. As a special case of coarsened or missing data, the corresponding estimators of *ξ*^*e*^ are shown to reach the *semiparametric efficiency bound* when both nuisance functions are specified correctly [20, 37].

Many machine learning (ML) approaches perform well by employing regularization to select among vast confounders. However, the induced *regularization bias* may propagate to invalidate the naive estimators of causal effects. The Neyman orthogonality motivated advancements such as debiased ML (DML) and target learning to remove the regularization bias, where one key component is deploying this Neyman-orthogonal score or EIF in (2) that is robust to the unknown patterns from confounders.

### 3.2 Causal Mediation Effect

Increasingly, investigators are interested in causal mediation analysis. For instance, upon evaluating the *total causal effect* of the exposure, they would like to further examine the *direct* or *indirect* pathways of the exposure, possibly through a *mediator* variable [35]. Following the literature [31, 32], we consider natural (pure) direct effects where the mediator is viewed as random in what follows.

For notation, a three-component (exposure *E*_*i*_, mediator *M*_*i*_, and outcome *Y*_*i*_) causal study design, where *M*_*i*_ ∈ Ω is the mediator from the exposure to an outcome. For instance, being exposed to cognitively stimulating activities *E*_*i*_ has been hypothesized to alter the cognitive functioning *Y*_*i*_ through the microbiome composition *M*_*i*_ as in our motivating example.

With *M*_*i*_ and *Y*_*i*_ the respective *observed* mediator and outcome, we define two counterfactual quantities under *e* ∈ {0, 1}:

- 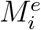 – *counterfactual mediator* for the exposure arm *E*_*i*_ = *e*;
- 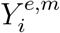 – *counterfactual outcome* for the *exposure-mediator combination* 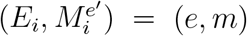.

In our example, the microbiome as a mediator can be affected by the cognitive stimulating activities and admits 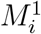 and 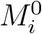, but for each subject *i*, only one of them can be observed, and hence, is counterfactual. The potential outcomes depend on both the mediator and exposure, which differ from the pure causal setting in (1) where the potential outcome only depends on exposure. The individual causal *indirect (mediation) effect* has been defined as 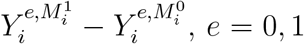, which answers a counterfactual question: what is the difference in the outcome when the value of mediator switches from 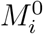 (under the control) to 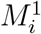 (under the exposure) while holding the exposure status at *e* [31].

In contrast, the individual causal *direct effect* is defined as 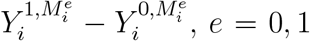. In our example, the direct stimulation effect on the subject *i* ‘s cognitive functioning while holding his or her microbiome composition constant at the level under no stimulation is 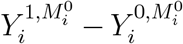. Further, if one assumes no interaction between *E*_*i*_ and *M*_*i*_, then the indirect and direct effects will no longer vary with *e*. Akin to the classical setting, those individual effects are unidentifiable due to their counterfactual nature.

In the presence of a mediator, we decompose the average causal effect (ACE) of *E* on *Y* in (1) as (drop the subscript *i*):

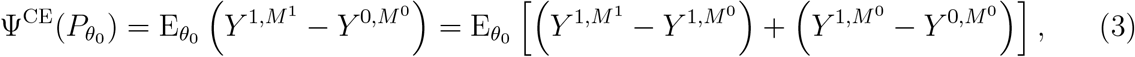

which respectively defines the *average mediation effect* (AME) and *average direct effect* (ADE):

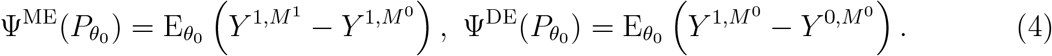

In particular, 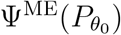 is the comparison of the potential outcome from the exposure arm when the mediator is switched on (*M* ^1^) and off (*M* ^0^). And 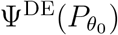 directly contrasts the potential outcomes between the two arms when the mediator was switched off (*M* ^0^) [35]. Both averaged across the entire population.

Not tied to any specific model, the definitions in (4) generalize previous discussions by **(author?)** [13] using linear structural equation model (SEM), where one posits two models, one for the outcome *Y* with 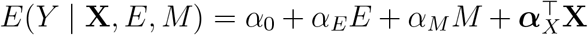, the other for the mediator *M* with 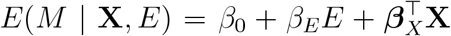. Hence, it is easily deduced from (4) that 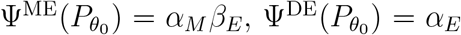. Often, it is of interest to test (1) the existence of mediation effect 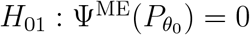, and (2) full vs. partial mediation by testing the direct effect 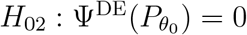.

#### 3.2.1 Assumptions

Denote by **X** ∈ *χ* a set of pre-exposure variables sufficient to account for selection bias, then under three stronger assumptions, 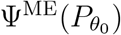 and 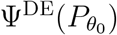 become identifiable.

D1. *Counterfactual Consistency* :

(i) The observed *M* = *EM* ^1^ + (1 − *E*)*M* ^0^ : if *E* = *e*, then *M*^*e*^ = *M*, w.p.1.

(ii) The observed *Y* = *EY* ^1,*m*^ + (1 − *E*)*Y* ^0,*m*^ : If *E*_*i*_ = *e* and the observed *M* = *m*, then *Y* ^*e*,*m*^ = *Y*, w.p.1.

D2. *Strong sequential ignorability* :

(i) *E* ⊥ {*Y* ^*e′*^, ^*m*^, *M*^*e*^} | **x** : given the observed confounders, the exposure assignment is statistically independent of the potential outcomes and potential mediators;

(ii) *Y* ^*e′*^,^*m*^ ⊥ *M*^*e*^ | *E* = *e*, **x** : given the observed exposure and confounders, the mediator is ignorable.

D3. *Positivity*: C3 holds and Pr(*M* | *e*, **x**) *>* 0, w.p.1. for each *m* ∈ Ω.

In D2, the two ignorability assumptions are sequential: first, given **x**, the exposure is assumed to be ignorable, which can sometimes be enforced by randomization [15]; second, the mediator is ignorable given the observed value of the ignorable exposure and **x**. It implies that among those subjects under the same exposure status and pre-exposure characteristics, the mediator can be regarded as if being randomized, which can be strong. Hence, we use sensitivity analysis to further validate the ignorability assumptions (see Section 4).

#### 3.2.2 Identification

Akin to the causal effect 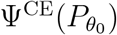, the causal direct and indirect effects can be constructed to be *multiply robust* and *locally efficient*, even with a large number of pre-exposure confounders. Under assumptions D1-D3, the *causal mediation functional* 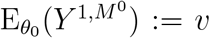 is identifiable [15, 31], which appeared in (4), we can further deduce that

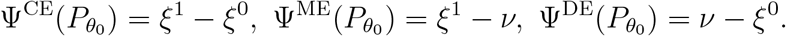

By the linearity, the EIF for the causal functional *ξ*^*e*^ in (2) can be used, and we only need to estimate the mediation functional *ν* for the estimation of AME and ADE. Using the Gautaex derivative and the strategy of a point mass [4], the efficient influence function (EIF) for *ν* is found to be [35]

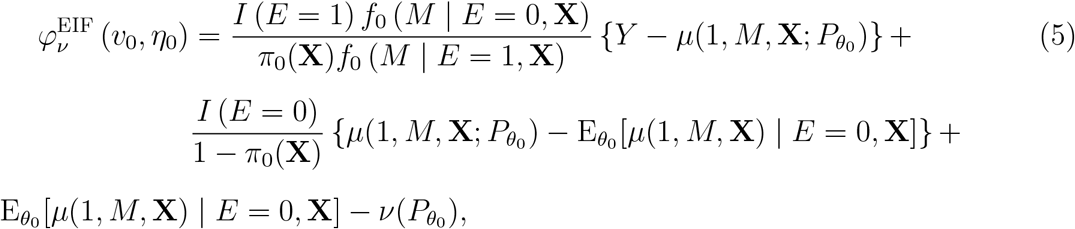

where *π*_0_(**X**_*i*_) is the true propensity score (PS) as in (2); also, the true outcome mean given exposure, mediator, and confounders is denoted as 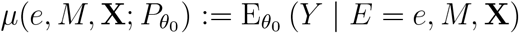. The additional part *f*_0_ (*M* | *E* = *e*, **X**) denotes the true conditional density of mediators; for discrete mediators, the probability mass function 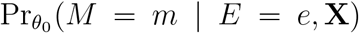 can be substituted.

To bypass estimating the density *f*_0_ (*M* | *E* = 0, **X**) when *M* is continuous, or even multidimensional, an alternative form has been proposed based on Bayes law [10], which substitutes the first component in (5) by

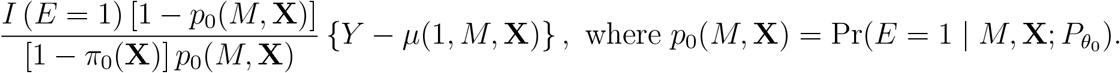

### 3.3 Confounding Selection

For the ubiquitous observational studies where randomization is not feasible or ethical, the utmost challenge is that the confounding mechanisms are unknown. To mitigate the potential bias, the investigators tend to collect as many baseline characteristics as possible, hoping to avoid unmeasured confounding. However, when the number of confounders approaches or exceeds the sample size, the model fitting becomes unstable without some filtering, either by adding a penalized term or using other strategies.

The previously introduced efficient influence functions (EIF) follow Neyman’s orthogonality condition [20]. Therefore, the resulting estimators should not be strongly affected by the quality or precision in estimating the confounding patterns [29]. In practice, different selection strategies may lead to distinct sets of confounders, hence propagating to yield different causal estimators. We briefly review some recent developments for selecting among the vast number of confounders in the context of causal mediation analyses.

#### 3.3.1 Double Machine Learning (medDML)

The debiased machine learning (DML) [4] is a popular approach deploying the orthogonal score (or EIF) and flexible nonparametric methods to handle the massive confounders. Conventional DML only estimates the causal effect 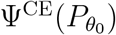. A recent development in [10] extends the scope into a causal mediation framework, which yields asymptotically normal and 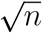-consistent average direct effect (ADE) and mediation effect (AME) estimators while automatically selecting among high-dimensional confounders.

Akin to the DML, two components are essential for obtaining well-behaved final estimators. First, to overcome the regularization bias in selecting the confounders, the Neyman orthogonal scores, or EIFs, are adopted, specifically, (5) for *ν* and (2) for *ξ*^*e*^. This procedure ensures the resulting estimators of ADE and AME are robust to misspecifications of the outcome and mediator models, also referred to as the “multiply robust” property.

Second, *cross-fitting* is another essential piece to prevent overfitting. By randomly splitting the sample and estimating the nuisance parameters in one half (*auxiliary* part) while estimating the ADE and AME in the other half (*main* part), one avoids potential overfitting in estimating the nuisance, which can occur, say, involving irrelevant confounders. Then, by swapping the role of main and auxiliary parts to obtain a second estimator, the full efficiency is recovered by taking their averages. Notably, it only requires weaker assumptions to shrink an empirical process term to zero, where the Donsker conditions do not apply in this high-dimensional setting [4].

The cross-fitting can be generalized to a *K*-fold version. To deploy the *K*-fold crossfitting procedure for estimating *v* and *ξ*^*e*^ in the mediation analyses, we denote the observed data by **O**_*i*_ = (*Y*_*i*_, *M*_*i*_, *D*_*i*_, *X*_*i*_)^⊤^ as an *i*.*i*.*d*. sample of size *n*. One first randomly partition the observation indices *I* = {1, …, *n*} into *K* disjoint subsets with equal size, denoted as *I*_*k*_ for *k* = 1, …, *K*, and denote by 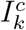 the complement set of *I*_*k*_.

Next, within each fold, some ML algorithm is applied to the observations in the complement set 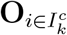 to estimate the nuisance parameters, which include the conditional outcome means, mediator densities, and propensity scores appeared in (5) and (2), collectively denoted as 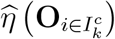. This process is repeated for all *K* folds, yielding *K* estimators by solving the empirical analog of EIFs (5) and (2) using estimating equations but with *η*_0_ replaced by 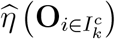 for each fold. The final estimators for *ν* and *ξ*^*e*^ are averaged across the *K* estimators.

Essentially, these two key components ensure the 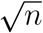-convergence of the estimated ADE and AME, where the regularity conditions are attained by various ML algorithms, including Lasso.

#### 3.3.2 Regularized Partial Correlation Network

The emergence of the *partial correlation network* modeling of psychosocial variables overcame the historical challenges of deciphering their complex interplay [9]. Further, regularization has been incorporated for a more interpretable and parsimonious network structure. Inherently differing from social networks that represent the connections among subjects, such networks reflect the connection (*edges*) between psychological variables (*nodes*) based on partial correlations [2, 24], also termed the *Gaussian graphical models* [17], which belong to the general class of *pairwise Markov random fields* [16, 27].

In the context of variable selection, consider a *p ×* 1 vector of potential confounders **X** (we suppressed the subscript *i* for each subject in what follows), denoted by *ρ*_*j*_ = Corr (*Y, X*_*j*_ | **X**_−*j*_) (*j* = 1, …, *p*) the *partial correlation coefficient* between the response *Y* and a confounder *X*_*j*_, after conditioning on other variables 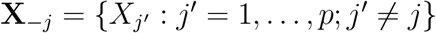 . Under the joint normal assumption, *ρ*_*j*_ captures the linear relationship between *Y* and *X*_*j*_, partial out other confounders. To estimate *ρ*_*j*_ from the sample, denote the *precision matrix* by 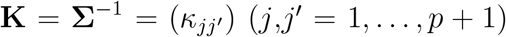 as the inverse of the variance-covariance matrix from (*Y*, **X**)^⊤^ ∼ *N* (**0, Σ**). It is readily shown that by standardizing the elements 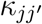, one obtains a *partial correlation matrix* 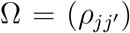, whose elements 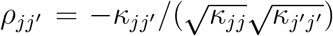. The first column of Ω (*j′* = 1) readily recovers each *ρ*_*j*_, the partial correlations between *Y* and *X*_*j*_.

However, during the estimation of **K**, one often encounters *spurious* edges that are not exactly zero due to the sampling variation [8]. To avoid overfitting, a *regularized* partial correlation network [12], or *partial correlation graphical Lasso* (glasso), has become prevalent. Accordingly, the log-likelihood adds a *L*_1_ penalty to the sum of absolute covariance values to yield the MLE of **K**:

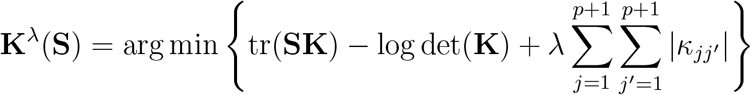

where **S** is the *sample* covariance matrix of (*Y*, **X**)^⊤^ and *λ* a tuning parameter to control the sparsity level. **K**^*λ*^(**S**) is well-suited for high-dimensional settings where *p* + 1 *> n*, under which det(**K**) = 0 and regularization is essential as in our context [39].

To choose *λ*, one can start by grid search and select the optimal one with minimal EBIC (Extended Bayesian Information Criterion) [3], which works well in retrieving the true network structure, especially when they are sparse. Let *l* denote the penalized loglikelihood, *m* the number of non-zero edges, and *n* the sample size, the EBIC is defined as −2*l* + *mlog*(*n*) + 4*γmlog*(*p* + 1), which contains a hyperparameter *γ* to control the preference over sparsity (i.e., fewer edges), suggested to be set between 0 and 0.5 [9], with higher values indicating more parsimonious network models are preferred.

Accordingly, the resulting penalized partial correlation coefficients 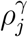 (range: [−1, 1]) encode the conditional association between two variables after controlling for all others and removing spurious connections. This is important for variable selection since *Y* and *X*_*j*_ being conditionally dependent is equivalent to 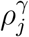 being non-zero, implying a stronger signal of *X*_*j*_,and hence a more *relevant* confounder after penalization. Thus, 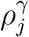 captures the relationship between the response and a relevant confounder, conditional on, or partial out, all other covariate effects, whereas 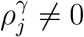 corresponds to irrelevant confounders, where *Y* and *X*_*j*_ are conditionally independent.

On the other hand, by directly penalizing elements in the variance-covariance matrix among the vast confounders, this variable selection strategy differs from conventional Lasso, which penalizes the coefficients in the regression model as in the DML approach. As the regularized partial correlations have become standard when estimating psychopathology networks, they provide a promising alternative for selecting confounders in the causal mediation analysis.

## 4 Real Data Analyses

### 4.1 Setup

With the motivating dataset described in Section 2, we now investigate whether exposure to loneliness and cognitively stimulating activities will alter cognitive functioning over time (after six months) via modulating the microbiome composition. This observational collected a large number of baseline characteristic variables, including the psychiatric and mental health instruments such as the PHQ9 severity score of depression, positive psychological traits such as Connor Davidson Resilience score, physical health such as sleep, and medications. They help mitigate unmeasured confounding but challenge our statistical inference since the number of confounders (*p* = 81) approaches the sample size (*n* = 92).

Hence, we need to select relevant confounders carefully. We first applied the regularized partial correlation network to screen the 81 potential confounders. Several hyperparameters *γ* in the EBIC were explored, from dense (*γ* = 0) to very sparse (*γ* = 0.5), each visualized by using the qgraph package in R [9]. Since *γ* influences the sparsity of the network connection, the chosen range covers a spectrum of complexities of the model.

The resulting network was estimated using the glasso algorithm, where we examined the number of edges and their strengths (i.e., the magnitude of partial correlations) across different *γ* values. We present the final models under *γ* = 0 and *γ* = 0.1 to compare the impact of the confounder selection. In the partial correlation network, the exposure of loneliness and cognitively stimulating activities yielded distinct sets of variables carrying non-zero coefficients, which were used as respective confounders for further causal mediation analyses.

We compared three procedures for deriving the causal mediation effects as follows:

(1) Traditional mediation analysis with network-selected confounders (*Non-EIF + Network-selection*) was conducted with the mediate() function from the mediation package. For the binary outcome, we deployed logistic regression for the outcome model and the linear model for the continuous mediator. For continuous outcome, we used the linear model for the mediator, and a generalized additive model (GAM) with smooth terms for the outcome to allow for a treatment-mediator interaction, which did not impose the same direct and indirect effect under treatment and control and hence is more flexible. Bootstrap was implemented with robust standard errors to account for potential heteroscedasticity.
(2) Double machine learning, or DML with network-selected confounders (*EIF + DML-selection + Network-selection*): the medDML() function from the causalweight package was applied using the selected confounders from the regularized partial correlation network. Cross-validation was employed to tune the hyperparameters.
(3) DML with full confounder set (*EIF + DML-selection*): the DML analysis was repeated using all 81 potential confounders, with the same configuration as in (2) but with an expanded set of confounders.

The three combinations were compared regarding their parameter estimates, standard error, and p-value for the average total, direct, and indirect effects under treatment and control.

Finally, we conducted sensitivity analyses to assess if the assumption of no unmeasured pre-treatment confounders holds using the function medsens() in the mediation package. This function yields a sensitivity parameter at which the causal indirect effect equals zero. More particular, it measures the correlation between the residuals from the mediator and outcome models, namely, let 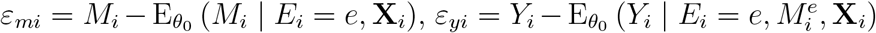, we have

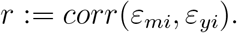

The interpretation is that when the assumption of no unmeasured pre-treatment confounders holds, the correlation between two model residuals should be close to zero. Thus, larger *r* values suggest less robust causal and mediation estimators, and caution is warranted [6].

We conducted the mediation analyses to investigate the two main scientific questions: first, the path from loneliness to microbiome to cognitive impairment, and second, the path from cognitively stimulating activities to microbiome to cognitive impairment. For each, we considered the continuous and binary outcome of cognitive impairment, measured by the MoCA instrument as a continuous score but also dichotomized by the clinical cutoff [28]. Sensitivities analyses for each analysis were presented by reporting the corresponding *r* values. Since this sensitivity analysis is implemented only for linear mediator and outcome models and linear mediator and binary probit outcome models, we assessed the sensitivity for binary MoCA outcome using the probit link even though the final results were based on the logit link.

### 4.2 Scientific Insights

#### 4.2.1 Loneliness → Microbiome → Cognitive impairment (MoCA)

With a prevalence rate of 20% to 35% among U.S. adults over the past decade [23], loneliness is considered the latest global health epidemic with serious health implications, including depression, cognitive impairment, hypertension, and frailty [14]. However, whether the impact of loneliness on cognitive impairment via the human microbiome, namely, the gut-brain axis, has not been previously investigated, our approach provided some insights as follows.

##### Binary MoCA outcome

(cognitive impairment vs. healthy control): When *γ* = 0 as shown in Table 2, the approach (2) deploying DML with network-selected confounders identified marginal indirect control effect (Ψ^ME^ = 0.106, *se* = 0.060, *p*-*val* = 0.079); while the DML with a full set of confounders in approach (3) identified an intensified effect (Ψ^ME^ = 0.141, *se* = 0.072, *p*-*val* = 0.050), suggesting being exposed to loneliness will increase the odds of being cognitively impaired, mediated through the microbiome alpha-diversity, in particular, the odds ratio of cognitive impairment and healthy control is *exp*(0.141) = 1.15 times when the values of alpha-diversity switches from 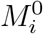 (under control) to 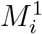 (under the exposure) while holding the actual exposure status at not exposing to loneliness. No significant effect appeared in approach (1).

##### Continuous MoCA outcome

No significant effects were identified for the continuous outcome.

#### 4.2.2 Cognitively Stimulating Activities → Microbiome → Cognitive impairment (MoCA)

The composite score of cognitive activity participation ranges from 1 to 5, with higher scores indicating more frequent participation in cognitively stimulating activities, which include education and training courses, reading, crossword puzzles, and playing chess or card games. Such activities have been shown to impact the cognitive functioning of older adults. For example, a study found that a 1-point increase in cognitive activity score was associated with a 33% reduction in the risk of Alzheimer’s disease (AD) [40]. Here, we validate the gut-brain axis to assess whether this path is partly through the microbiome using causal mediation analyses.

##### Binary MoCA outcome

When *γ* = 0.1 as shown in Table 3, the approach (2) using DML with network-selected confounders identified a significant indirect exposure effect (Ψ^ME^ = 0.014, *se* = 0.005, *p*-*val* = 0.013), supporting the mediation effect through the microbiome alpha-diversity, in particular, the odds ratio of cognitive impairment and healthy control is *exp*(0.014) = 1.014 times when the values of alpha-diversity are switch on (i.e., 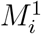 under the exposure) while holding the actual exposure to cognitive stimulates.

##### Continuous MoCA outcome (the higher, the less cognitively impaired)

With *γ* = 0.1, the partial network selected only LOT-R Total Score (Optimism) and 3D Wisdom Scale - Cognitive dimension as confounders. The mediation package with these two confounders in approach (1) identified a significant total effect (Ψ^TE^ = 1.941, *se* = 0.853, *p*-*val* = 0.024), as well as direct effect under the exposure and control (Ψ^DE^ = 1.909, exposure *p*-*val* = 0.024, control *p*-*val* = 0.018); they suggest that being exposed to cognitive activities can improve the cognitive functioning.

Interestingly, the network selected three more confounders with less stringent EBIC where *γ* = 0, including Hollingshead Index of Social Position (ISP) - Current Status, Neff Self-Compassion Scale score, and Nutrition Screening Checklist Total Score. The previous two effects were no longer significant after adding them.

Also, the DML approach with network-selected confounders in (2) did not find any significant effect for *γ* = 0 or 0.1.

The DML approach in (3) with a full set of confounders identified a significant total effect (Ψ^TE^ = 1.997, *se* = 0.971, *p*-*val* = 0.040), yet the direct effects were no longer significant.

#### 4.3 Methodological Implications

By comparing the analyses across the three combinations in selecting confounders and mediation estimation, we present some implications in statistical methods as follows.

##### In network-based confounder selection, the confounding sets were stable for the exposure of loneliness across different γ values, which induces more comparable final causal estimators

For example, when *γ* = 0, this loose EBIC criteria only yields one additional confounder of the SF-36 Mental Component Scale, compared with more stringent *γ* = 0.1 or 0.2, which both selected 12 confounders. It partly explained the similar final results of loneliness to microbiome to binary cognitive impairment (MoCA) when comparing *γ* = 0 and 0.1. However, this was not the case for cognitively stimulating activities, where the selected sets were much smaller. For example, when *γ* = 0, five confounders were selected, which reduced to two when *γ* = 0.1 and one when *γ* = 0.2, as shown in Table 1.

**Table 1:** Non-zero Connections to Loneliness (uclalst) and Cognitive Stimulus (csascore) with Varying *γ* Values for Sparsity.

| **Treatment** | **Confounder** | $\gamma = 0$ | $\gamma = 0.1$ | $\gamma = 0.2$ | $\gamma \geq 0.24$ |
| --- | --- | --- | --- | --- | --- |
| Loneliness | mencomp | -0.011 | – | – |  |
|  | ldr2 | 0.080 | 0.065 | 0.049 |  |
|  | lotrt | -0.067 | -0.065 | -0.060 |  |
|  | mlq_ps | -0.030 | -0.034 | -0.035 |  |
|  | cse_sff | -0.056 | -0.062 | -0.063 |  |
|  | nefftot | -0.129 | -0.127 | -0.123 |  |
|  | wsdm_cd | 0.080 | 0.062 | 0.042 | No connection |
|  | sdw_sa | -0.013 | -0.009 | -0.001 |  |
|  | sdw_psb | -0.030 | -0.020 | -0.007 |  |
|  | phycomp | -0.081 | -0.072 | -0.060 |  |
|  | prsd8a_ss | 0.099 | 0.072 | 0.045 |  |
|  | prsi_ss | 0.105 | 0.084 | 0.059 |  |
|  | prdsa_ss | -0.090 | -0.069 | -0.048 |  |
| Cognitive Stimulus | socposc | -0.036 | – | – |  |
|  | lotrt | -0.078 | -0.046 | -0.006 |  |
|  | nefftot | -0.028 | – | – | No connection |
|  | wsdm_cd | 0.054 | 0.012 | – |  |
|  | nsictot | -0.014 | – | – |  |

**Table 2:**
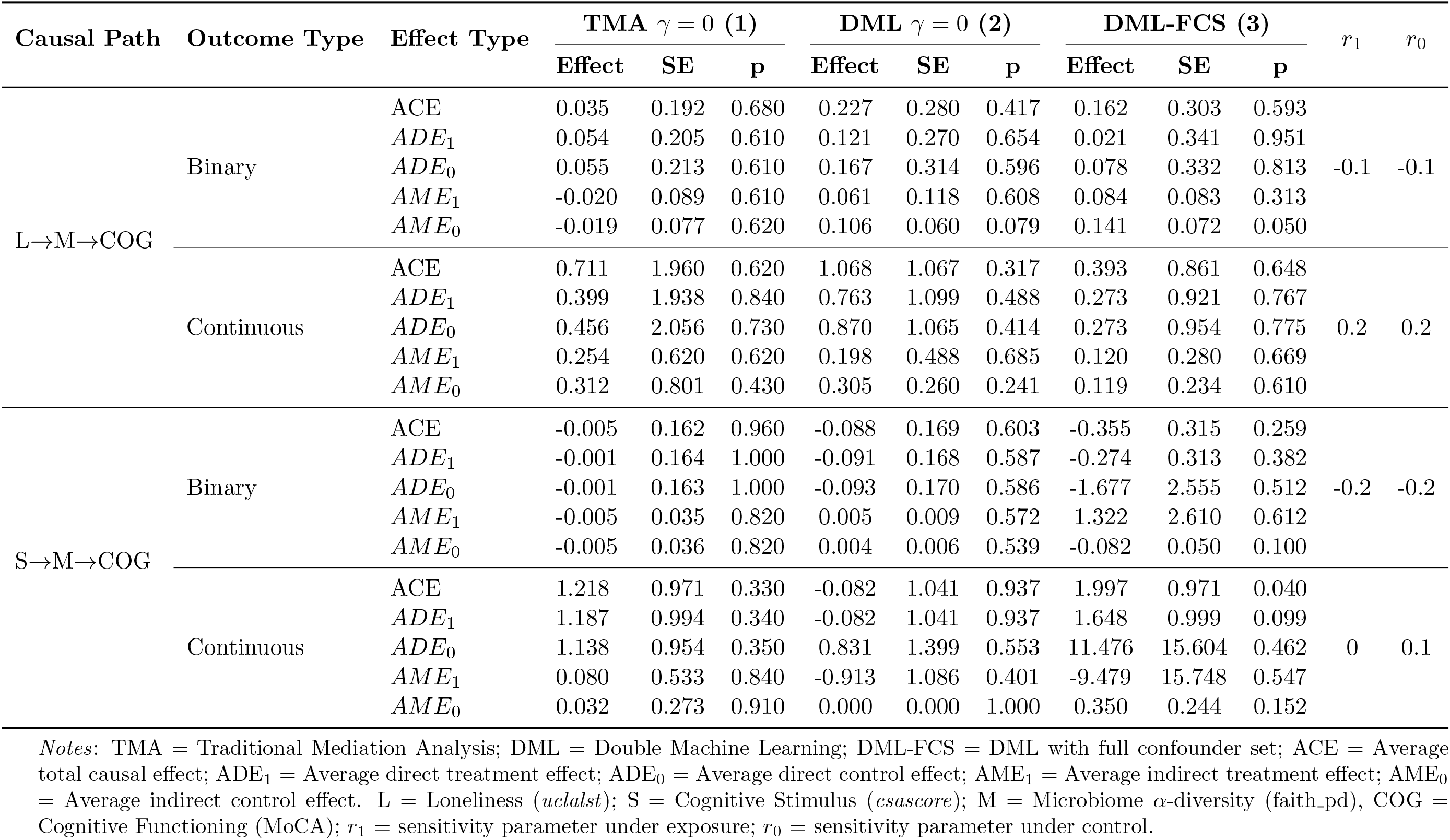
Mediation Effects of Loneliness/Cognitive Stimulus on Cognitive Functioning via Gut Microbiome (*γ* = 0)

**Table 3:**
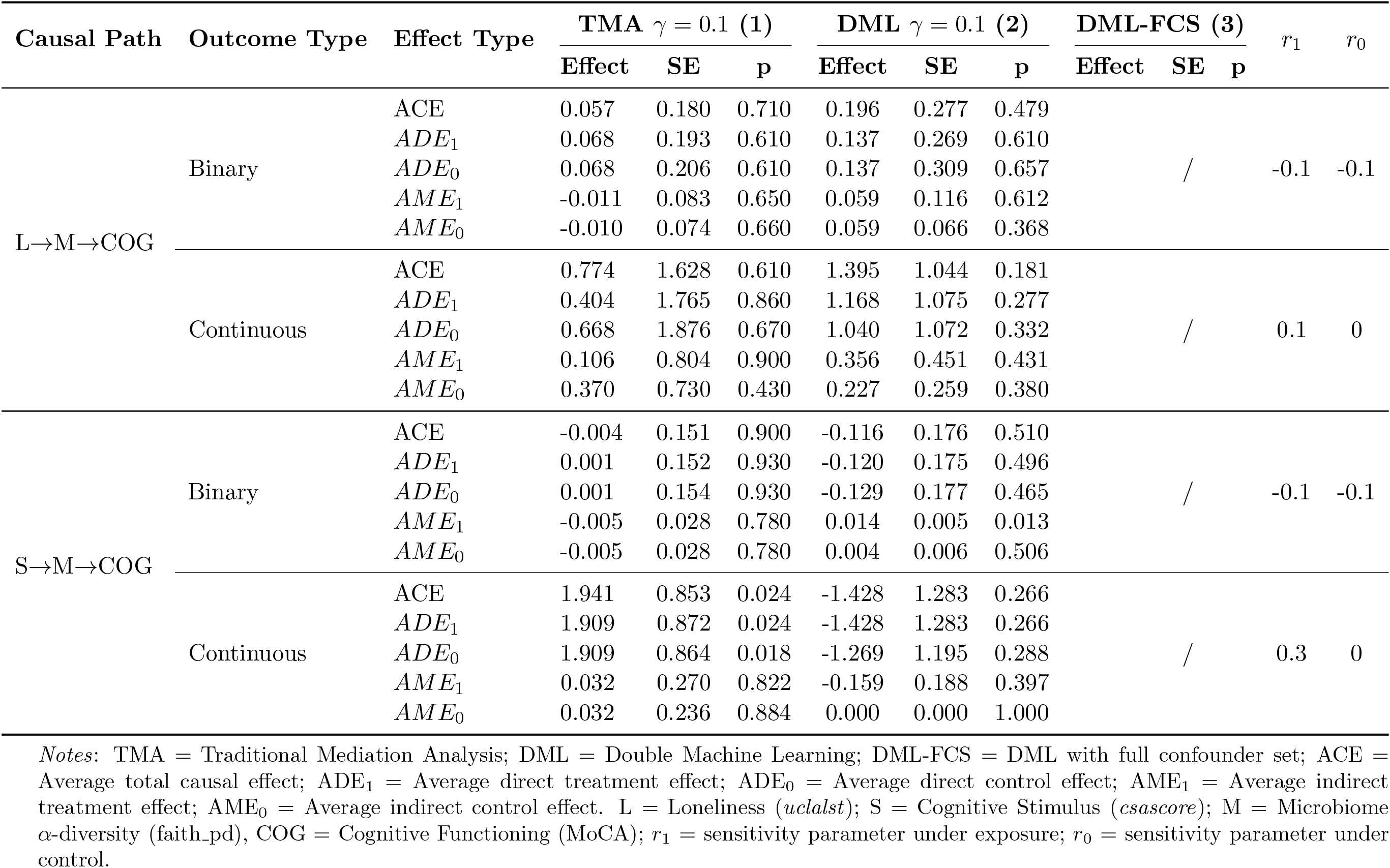
Mediation Effects of Loneliness/Cognitive Stimulus on Cognitive Functioning via Gut Microbiome (*γ* = 0.1)

##### DML with distinct confounder sets can lead to different mediation estimations, regardless of continuous or binary outcomes

For example, when *γ* = 0, albeit the same sign for the indirect control effect of microbiome under approaches (2) and (3), DML with the full sets yielded a larger effect (0.106 *vs*. 0.141) and a smaller p-value (0.079 *vs*. 0.050) from loneliness to the binary MoCA. This is especially the case if more distinct confounder sets are selected. In particular, for the path of cognitively stimulating activities to microbiome to binary MoCA, the indirect exposure effect was significantly positive when *γ* = 0.1 but almost negligible when *γ* = 0 (0.014 *vs*. 0.005).

##### DML, in general, is more sensitive to detecting total, direct, or indirect effects than traditional mediation analyses, even with the pre-selected subsets of confounders

Across the eight sets of evaluated hypotheses, the DML-based approach detected significant or marginally significant effects in six, where four were from the DML with the full confounder set. This observation highlights that the automatic screening of confounders in DML is already quite powerful in mitigating the overfitting and deducing consistent causal effects, thanks to the implemented cross-fitting and Neyman orthogonality.

##### DML seemed more effective in detecting any effect in binary than continuous outcomes

Four of the six significant effects detected by the DML-based approach were on the binary outcomes, where the corresponding continuous outcome did not yield significant results. For example, in the path of cognitive stimulating activities to microbiome to binary MoCA, DML with network-based confounder detected a significant indirect exposure effect (Ψ^ME^ = 0.014, *se* = 0.005, *p*-*val* = 0.013) while no strong effect was identified in continuous the MoCA (*p*-*val* = 0.397).

##### The network-based confounder selection can help add to the DML approach, especially in improving the robustness and stability of the model fit

For instance, in the path of cognitively stimulating activities to microbiome to continuous MoCA, DML with the full sets in approach (3) yielded inflated direct and indirect effects compared with approach (2) that deploys the network-selected sets (*e*.*g*., 0.83 *vs*. 11.48), which could be an artifact of the sample-splitting over a small sample size.

## 5 Discussion

In this paper, we addressed a timely issue of quantifying the causal mediation effect encountering high-dimensional confounders. Under the counterfactual framework, we first showed that the average causal effect (ACE) is decomposed into the average indirect (or mediation, AME) and direct effects (ADE), which facilitated constructing the nonparametric target causal functionals without attaching to any specific model. Later, two confounding selection strategies were carefully studied, including double machine learning (DML) and regularized partial correlation network. To our knowledge, these two promising approaches have not been compared in the growing causal mediation setting under high dimensionality.

We, hence, offered thorough comparisons among various combinations to evaluate their impacts on the final estimation of target parameters, which not only guides real-world applications for practitioners but also incentivizes future advancements for this important topic. In our motivating data from a longitudinal observational study on the human microbiome, we encountered high dimensionality in both the mediator and confounders, coupled with a small to moderate sample size. For the mediator of microbiome taxa counts, we leveraged the feature aggregation to enrich signals and domain-specific structures using diversity metrics [22], which have been well-recognized in the field [7, 25]. For the massive confounders, we considered the two confounding selection strategies. Along with the efficient influence functions for the causal mediation effects, three combinations were carefully studied to demystify causality in the “gut-brain axis.” Our results are consistent with the scientific literature but offer nuanced methodological implications on how the confounding selection impacts the final causal target parameter estimation, above and beyond the real-world scientific insights.

However, the study results are still limited by the relatively small sample size and the imputation of the missing covariates in the follow-up visit. Accordingly, we also discussed the possibility of using the two strategies in conjunction to improve the stability of model fit. Nonetheless, the sensitivity analyses confirmed that the crucial assumption of no unmeasured pre-treatment confounders holds in the various hypotheses that we tested.

Finally, our causal mediation analyses showcased the exposure impact of loneliness and cognitively stimulative activities on cognitive functioning for the aging population, which is mediated by their microbiome composition, supporting the potential for gut microbiota targets in preventing or treating cognitive decline. Those thorough comparison studies are valuable in many other growing fields encountering high dimensionality, such as the metabolomics or functional connectivity in neuroimage that are commonly hypothesized as the mediator [1, 18]).

In summary, our results highlighted the practicality and necessity of the discussed methods in mitigating selection bias in causal mediation analysis, especially when the dimension of mediator and confounders exceed the sample size.

## A Appendix

